# Spider web DNA: a new spin on noninvasive genetics of predator and prey

**DOI:** 10.1101/011775

**Authors:** Charles C. Y. Xu, Ivy J. Yen, Dean Bowman, Cameron R. Turner

## Abstract

Noninvasive genetic approaches enable biomonitoring without the need to directly observe or disturb target organisms. Environmental DNA (eDNA) methods have recently extended this approach by assaying genetic material within bulk environmental samples without *a priori* knowledge about the presence of target biological material. This paper describes a novel and promising source of noninvasive spider DNA and insect eDNA from spider webs. Using black widow spiders (*Latrodectus* spp.) fed with house crickets (*Acheta domesticus*), we successfully extracted and amplified mitochondrial DNA sequences of both spider and prey from spider web. Detectability of spider DNA did not differ between assays with amplicon sizes from 135 to 497 base pairs. Spider DNA and prey eDNA remained detectable at least 88 days after living organisms were no longer present on the web. Spider web DNA may be an important tool in conservation research, pest management, biogeography studies, and biodiversity assessments.

## Introduction

As dominant predators of arthropod communities in natural and agricultural ecosystems, spiders are important ecological indicators that reflect habitat quality and change across trophic levels (Churchill 1997; Clausen 1986). Monitoring the species diversity and abundance of spider assemblages facilitates natural resource management (Pearce and Venier 2006). Spiders are enormously diverse (~ 44,000 described species; Platnick 2013) and difficult to identify. Morphological identification of spiders relies primarily on differences in copulatory organs (Huber 2004) and many complications can prevent identification such as the inability to identify juveniles, extreme sexual dimorphism, size differences between life stages, and genital polymorphisms (Barrett and Hebert 2005; Brennan *et al.* 2004; Huber and Gonzalez 2001). Other major issues include the ever decreasing availability of expertise necessary for traditional taxonomy as well as the significant training required to learn taxonomic skills (Hopkins and Freckleton 2002). In the face of such challenges to morphological taxonomy, genetic identification methods are growing in popularity because of decreasing costs and ease of use. DNA barcoding, the use of a short and standardized fragment of DNA to identify organisms, has gained significant traction within the last decade (Jinbo *et al.* 2011). In particular, the use of DNA barcodes for species identity and systematics of spiders has proven successful in multiple studies (Astrin *et al.* 2006; Barrett and Hebert 2005; Robinson *et al.* 2009). The most commonly used genetic marker is the cytochrome oxidase subunit I (COI) mitochondrial gene because of its designation as the standard DNA barcode (Hebert *et al.* 2003). Mitochondrial markers are also ideal for detecting low quantity and quality DNA from environmental or gut samples because each cell contains hundreds to thousands of mitochondrial genomes (Hoy 1994) and there is a positive correlation between gene copy number and detection success (Agustí *et al.* 2003b; Chen *et al.* 2000).

Spiders have a great diversity of life histories and various sampling methods are employed in capturing them including vacuum sampling, sweep netting, pitfall traps, and visual searches. Experiments testing the efficacy of traditional spider sampling methods show high variability between methods as well as inconsistency across spatial and temporal scales (Churchill and Arthur 1999; Green 1999; Merrett and Snazell 1983). Sampling duration is also an important factor as short-term sampling has been found to reduce the number of recorded species by up to 50% (Riecken 1999). In this paper, we propose a new biomonitoring tool that would complement existing methods: DNA from spider web. While spider web has been found to effectively collect pollen, fungal spores and agrochemical sprays (Eggs and Sanders 2013; Samu *et al.* 1992), no study, to our knowledge, has assessed spider web as a potential source of genetic material. We hypothesized that spider web could simultaneously provide a noninvasive genetic sample (spider DNA) and an environmental DNA sample (prey DNA). Noninvasive genetic sampling uses extraorganismal material like feces, hair, and feathers from individual organisms for genetic analysis without the need to contact target organisms (Beja-Pereira *et al.* 2009). Environmental DNA (eDNA) sampling uses genetic material from environmental mixtures like water or soil without isolating target organisms or their parts (Turner *et al.* 2014).

Although noninvasive genetic sampling is most common for vertebrates, it has been successfully applied to arthropod exuviae and frass (Feinstein 2004; Petersen *et al.* 2006). Webs are an abundant and easily collected spider secretion that may provide spider DNA. Spider webs may also contain eDNA from captured prey and other local organisms, functioning as natural biodiversity samplers. This idea parallels recent molecular studies using mosquitos, ticks, leeches, and carrion flies to sample local animal biodiversity (Calvignac-Spencer *et al.* 2013, Gariepy *et al.* 2012, Schnell *et al.* 2012, Townzen *et al.* 2008). Previous studies have successfully used mitochondrial DNA markers to detect spider prey from gut contents, but this requires physically capturing and killing spiders (Agustí *et al.* 2003a; Sheppard *et al.* 2005). Furthermore, traditional taxonomic identification of spider prey items is time-consuming, subject to human error, and accurate only to the order level (Salomon 2011). Spider webs may provide a unique noninvasive opportunity to study arthropod communities without the need to directly observe spider or insect.

Here, we tested the feasibility of extracting, amplifying and sequencing DNA of black widow spiders, *Latrodectus* spp. (Araneae: Theridiidae), and their prey, the house cricket *Acheta domesticus* (Orthoptera: Gryllidae), from black widow spider webs. Because extraorganismal DNA in spider webs is exposed to environmental degradation and may exist in short fragments, we used nested primer sets to test the effect of amplicon size on detection probability.

## Materials and methods

### Web collection

The black widow spider exhibit at the Potawatomi Zoo in South Bend, Indiana was inhabited by a single female western black widow spider (*Latrodectus hesperus*) before its death on November 19, 2011. The spider was fed 2 medium sized house crickets (*A. domesticus*), on a weekly basis by zookeepers. The exhibit measured 40 cm by 40 cm by 40 cm and contained a few twigs, a small piece of wood, and wood shavings lining its floor. 88 days after the death of the spider, a web sample was collected from the exhibit on February 15, 2012, which will be referred to as “Lhes_zoo”. The duration of inhabitance within the exhibit prior to the sample collection date is unknown. Three individual enclosures measuring 35 cm by 30 cm by 35 cm were constructed with plywood and acrylic sheeting. All enclosures were decontaminated with 10% bleach and installed at the Potawatomi Zoo in South Bend, Indiana.

Three female southern black widow spiders (*Latrodectus mactans*) were purchased from Tarantula Spiders (http://tarantulaspiders.com/). The spiders were hatched from egg sacs collected in Marion County, Florida, USA and raised on 2-3 housefly maggots (*Musca domestica*) twice per week before delivery to the Potawatomi Zoo. A single live *L. mactans* and a decontaminated branch for web building were placed into each enclosure on April 26, 2012 (Figure 1). Each *L. mactans* was immediately fed two medium-sized crickets by placing them onto web. Web samples were collected from each enclosure 11 days later on May 7, 2012, which will be referred to as “Lmac_1”, “Lmac_2”, and “Lmac_3”. All web samples were collected by twisting single-use, sterile plastic applicators to spool silk strands. No organism body parts or exuviae were visible in any web samples but cricket parts and spider feces were clearly evident on the bottom of the enclosures. Applicator tips were snipped into 1.5 mL microcentrifuge tubes using 10% bleach decontaminated scissors before storing at -20°C.

**Figure 1.**
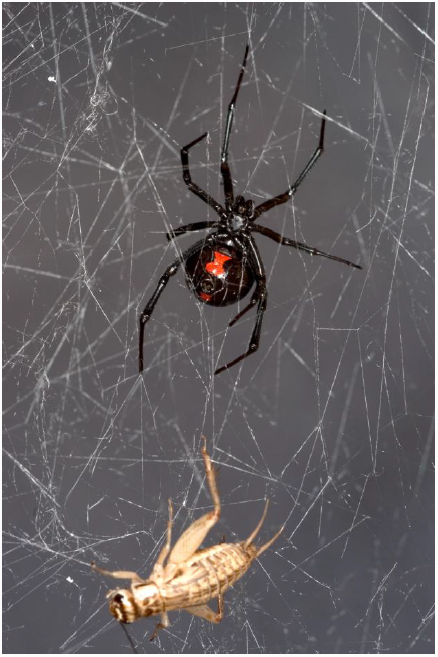
Southern black widow spider (*Latrodectus mactans*) with its prey house cricket (*Acheta domesticus*) trapped in spider web.

### DNA extraction

DNA extractions from web samples were conducted using a modified extraction protocol for shed reptile skins (Fetzner 1999). One negative control containing no web was also extracted. 800 *μ*L of cell lysis buffer (10 mM Tris, 10 mM EDTA, 2% sodium dodecyl sulfate [SDS], pH 8.0) and 8*μ*L of proteinase K (20 mg/L) were added to 1.5 mL microcentrifuge tubes containing web samples followed by 10-20 inversions and incubation at 55°C for 4 hours. Upon reaching room temperature, 4 *μ*L of RNase A (10 mg/mL) were added to each sample followed by 20 inversions. Samples were incubated at 37°C for 15 min and then brought back to room temperature. 300 *μ*L of protein precipitation solution (7.5 M ammonium acetate) were added to each sample and vortexed for 20 seconds followed by incubation on ice for 15 min. Samples were then centrifuged at 16,873 rcf for 3 min. Supernatants were transferred to new 2 mL microcentrifuge tubes containing 750 *μ*L of ice cold isopropanol and inverted 50 times before centrifugation at 14,000 rpm for 2 min. All supernatants were drained and 750 *μ*L of 70% ethanol was added to each sample followed by centrifugation at 14,000 rpm for 3 min. All liquids were removed and samples were air dried. DNA pellets were rehydrated using 100 *μ*L of low TE buffer (10 mM Tris, 0.1 mM EDTA).

### Primer design

To detect *Latrodectus* DNA, we designed four nested primer sets based on an alignment of *Latrodectus* COI DNA barcoding sequences obtained from the National Center for Biotechnology Information (NCBI) GenBank database. All four assays included the same forward primer but different reverse primers, producing amplicons of 135 bp, 257 bp, 311 bp, and 497 bp respectively (Table 1). To detect prey DNA, we designed a set of primers that specifically targets the DNA barcoding region of the COI gene in *A. domesticus*, which produces an amplicon of 248 bp (Table 1).

**Table 1:**
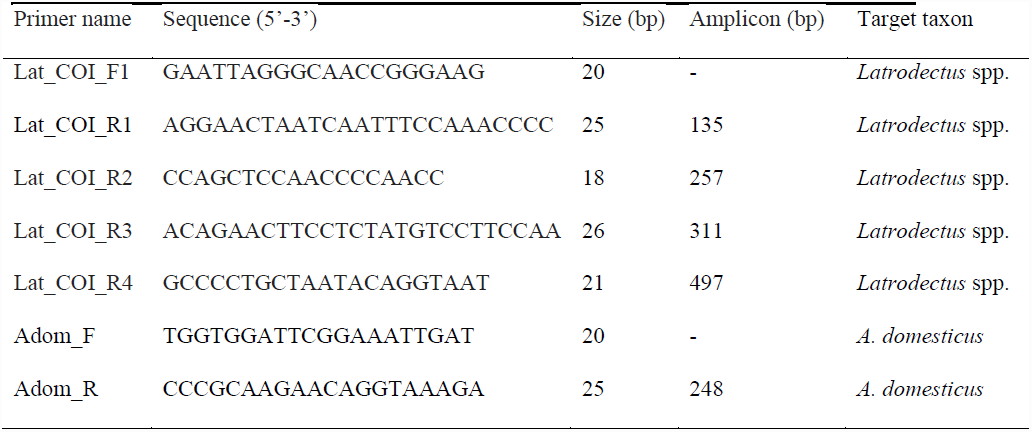
PCR primers designed to amplify the DNA barcoding region of the cytochrome oxidase subunit I gene of target species. All *Latrodectus* spp. primer sets are nested and use the same forward primer.

### DNA amplification

All DNA samples were amplified in polymerase chain reactions (PCR) of 20 *μ*L containing 13.28 *μ*L of ddH_2_O, 2 *μ*L of 5 PRIME® 10x Taq Buffer advanced, 2 *μ*L of 5 PRIME® Magnesium Solution at 25 mM, 0.4 *μ*L of dNTP at 2.5 mM, 0.12 *μ*L of 5 PRIME® Taq DNA polymerase at 5 U/*μ*L, 0.6 *μ*L of forward and reverse primers at 10 *μ*M, and 1.0 *μ*L of DNA template using Eppendorf Mastercycler® pro thermocyclers. Cycling conditions were as follows: 94°C/5 min, 55X (94°C/20 s, 54.4°C/35 s, 72° C/30 s), 72° C/7 m, 4° C/hold. Each *Latrodectus* spp. primer set was used to amplify all DNA samples with 10 technical replicates to measure detection probability for different amplicon sizes. All DNA samples were amplified with 2 technical replicates using the *A. domesticus* primer set. Negative control reactions to detect contamination were included in every batch. Gel electrophoresis was conducted using 5 *μ*L of PCR product mixed with 3 *μ*L of loading dye and 10 *μ*L of ddH_2_O. Multiple wells were loaded with 5 *μ*L of 100 bp ladder (Promega) on each gel. Technical replicates showing amplicons of the expected size were pooled and purified using ExoSAP-IT (Affymetrix). Sanger sequencing using ABI BigDye chemistry (Life Technologies) was conducted on an ABI 3730xl 96-capillary sequencer by the University of Notre Dame Genomics Core Facility. Sequencing chromatograms were primer- and quality-trimmed in Sequencher (ver. 5.0; Gene Codes Corp.). BLASTn searches of the NCBI GenBank database (http://www.ncbi.nlm.nih.gov; Benson *et al.* 2012) were used for taxonomic identification of COI barcode sequences.

## Results

All extraction and PCR negative controls produced no amplification. Using the nested primer sets, we successfully amplified 135 bp, 257 bp, 311 bp, and 497 bp of *Latrodectus* spp. COI from web DNA samples (Figure 2). With the exception of zero amplification for the 265 bp PCR assay from two samples, 2-10 technical replicates of each PCR assay successfully amplified from all samples. DNA sequences obtained from enclosure samples, “Lmac_1”, “Lmac_2”, and “Lmac_3”, were confirmed by NCBI BLAST to be *L. mactans* and DNA from the zoo exhibit sample, “Lhes_zoo”, was confirmed to be *L. hesperus*. Amplicon size had no effect on PCR success based on the number of successful PCR replicates (ANOVA, F = 1.941, d.f. = 3, *P* = 0.194). We also successfully amplified 248 bp of *Acheta domesticus* COI from eDNA samples. Both PCR duplicates from all four web samples were positive and all resulting DNA sequences were confirmed by NCBI BLAST to be *A. domesticus*. The zoo exhibit web sample, “Lhes_zoo”, was collected 88 days after the death and removal of both spider and prey, demonstrating substantial persistence of web DNA. All DNA sequences generated in this study are provided in Table S1 (Supplementary Data).

**Figure 2.**
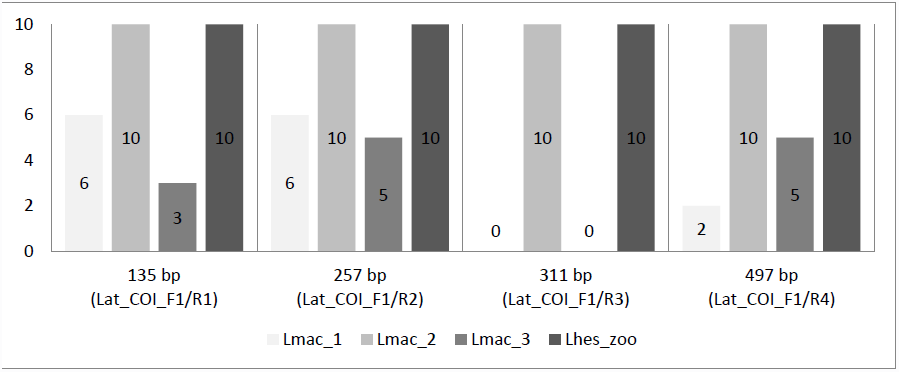
Success in detecting the mtDNA cytochrome c oxidase subunit I (COI) locus of *Latrodectus* spp. from web samples as measured by the number of positive PCRs out of 10 replicates. Samples “Lmac_1”, “Lmac_2”, and “Lmac_3” were tested for *Latrodectus mactans* while “Lhes_zoo” was tested for *Latrodectus hesperus* using the same nested “Lat_COI” primer sets.

## Discussion

The present study represents, to our knowledge, the first demonstration of spider web as a source of noninvasive genetic material. Spider web is an ideal source of noninvasive genetic material for spiders because web can be found and collected without direct observation of target organisms. Unlike most spiders, which are small, mobile, and elusive, webs are relatively large, stationary, and usually clearly visible, making sample collection more efficient. Spider webs may also remain after the inhabitant moves or dies, which increases detection probability, especially for the more elusive spider species. Webs can also exist in great abundance. For example, web coverage may reach up to more than 50% of land area in agricultural fields (Sunderland *et al.* 1986). Spider webs have already been utilized by citizen scientists to assess spider biodiversity through visual analysis of web structure (Gollan *et al.* 2010). It could be possible to implement similar citizen science initiatives to collect web samples for DNA analysis.

We hypothesize that spider web DNA originates either from microscopic pieces of fecal matter, setae, and exuviae adhered to silk strands or directly from the silk gland exudate, which may contain cells and mitochondria shed from silk glands. Because black widow spiders are orb weavers that generate large three-dimensional cobwebs consisting of sheets dotted with glue droplets (Zevenbergen *et al.* 2008), they were ideal to use in this experiment. Certain black widow spiders like *L. mactans* and *L. hesperus* are common venomous pests so spider web DNA could be a particularly useful tool for pest surveillance (Lewitus 1935). Because webs are easier to find and collect than live spiders, spider web DNA could also help monitor low density populations and determine invasive fronts of invasive widow spiders such as the brown widow, *Latrodectus geometricus*, in southern California and the Australian redback, *Latrodectus hasseltii*, in New Zealand and Japan (Vetter *et al.* 2012; Vink *et al.* 2011). Besides pests and invasives, many spider species like the red katipo (*Latrodectus katipo*) are threatened or endangered and hundreds if not thousands more are listed as “Data Deficient” but are probably at risk of decline (Sirvid *et al.* 2012). Spider web DNA could be particularly useful in easily providing occurrence and genetic diversity data for these rare species of concern. As a noninvasive biomonitoring method, spider web DNA could be used for conservation and taxonomy without sacrificing organisms that are already threatened by human disturbance. The collection and genetic analysis of spider webs could also serve spider biogeography studies, which require large-scale sampling across wide geographic ranges (Garb *et al.* 2004). Even silk from organisms that do not weave webs such as tarantulas and moth larvae may still yield viable DNA, but further experimentation is needed. This may be applicable towards molecular studies of trapdoor spiders, which construct burrows using silk but are extraordinarily difficult to capture for genetic sampling (Cooper *et al.* 2011).

Although the efficacy of spider web eDNA needs to be validated with samples from the field, this is the first demonstration that DNA of other insects can be extracted from spider webs. Spider predation can serve as a useful proxy to monitor local arthropod biodiversity. In some environments such as temperate forests, approximately 40% of arthropod biomass is annually consumed by spiders (Moulder and Reichle 1972). Although spider predation cannot be concluded from the mere presence of DNA on spider web, it does indicate the local proximity of those organisms. The ability to target particular species could be useful in monitoring low density populations of pest, invasive, or endangered insects. Future work using massively parallel sequencing on spider web eDNA could reveal entire assemblages of arthropods in a cost-effective manner, especially with the rapid advancement and decreasing costs of such technologies (Shokralla *et al.* 2012). Spider web eDNA may complement traditional assessment methods of local arthropod biodiversity and potentially reveal previously undiscovered biodiversity through improved sensitivity and sampling effort (Neilsen and Laurence 2000). Such information regarding species diversity is critically important in conservation planning and environmental impact assessments (Kremen *et al.* 1993, Rosenberg *et al.* 1986).

In conclusion, we provide the first demonstration that noninvasive DNA of spider and its prey can be extracted from spider web and be used to identify organisms to species. This method is low-cost, efficient, and does not require significant taxonomic expertise. Spider web DNA is a promising tool for the biomonitoring of spiders and other arthropods, especially if combined with the power of massively parallel sequencing.

## Author Contributions

CCYX and CRT designed the research. CCYX, CRT, IJY, and DB performed the research. CCYX and CRT wrote the paper with help from IJY and DB.

## Acknowledgements

We thank Larry Richey for his help in constructing the enclosures and we are indebted to Scott Camazine for sharing his photograph for Figure 1. We would also like to thank the Potawatomi Zoo for their cooperation as well as all the members of the Lodge lab and especially Dr. David Lodge for their support of this project.

## Data Accessibility

All DNA sequences generated in this study are provided in Table S1 (Supplementary Data) and will be archived in NCBI Genbank before publication of this manuscript.

## References

Agustí N, Shayler SP, Harwood JD, Vaughan IP, Sunderland KD, Symondson WOC (2003a) Collembola as alternative prey sustaining spiders in arable ecosystems: prey detection within predators using molecular markers. Molecular Ecology, 12, 3467–3475.

Agustí N, Unruh TR, Welter SC (2003b) Detecting *Cacopsylla pyricola* (Hemiptera: Psyllidae) in predator guts using COI mitochondrial markers. Bulletin of Entomological Research, 93, 179–185.

Astrin JJ, Huber BA, Misof B, Klütsch CFC (2006) Molecular taxonomy in pholcid spiders (Pholcidae, Araneae): evaluation of species identification methods using CO1 and 16S rRNA. Zoologica Scripta, 35, 441–457.

Barrett RDH and Hebert PDN (2005) Identifying spiders through DNA barcodes. Canadian Journal of Zoology, 83, 481–491.

Beja-Pereira A, Oliveira R, Alves, PC, Schwartz MK, Luikart G (2009) Advancing ecological understandings through technological transformations in noninvasive genetics. Molecular Ecology Resources, 9, 28–36.

Benson DA, Karsch-Mizrachi I, Clark K, Lipman DJ, Ostell J, Sayers EW (2012) GenBank. Nucleic Acids Research, 40, D48–D53.

Brennan KEC, Moir ML, Majer JD (2004) Exhaustive sampling in a Southern Hemisphere global biodiversity hotspot: inventorying species richness and assessing endemicity of the little known jarrah forest spiders. Pacific Conservation Biology, 10, 241–260.

Calvignac-Spencer S, Merkel K, Kutzner N, Kühl H, Boesch C, Kappeler PM, Metzger S, Schubert G, Leendertz FH (2013) Carrion fly-derived DNA as a tool for comprehensive and cost-effective assessment of mammalian biodiversity. Molecular Ecology, 22, 915–924.

Chen Y, Giles KL, Payton ME, Greenstone MH (2000) Identifying key cereal aphid predators by molecular gut analysis. Molecular Ecology, 9, 1887–1898.

Churchill TB (1997) Spiders as ecological indicators: an overview for Australia. Memoirs of the National Museum of Victoria, 56, 331–337.

Churchill TB, Arthur JM (1999) Measuring spider richness: effects of different sampling and spatial and temporal scales. Journal of Insect Conservation, 3, 287–295.

Clausen HIS (1986) The use of spiders (Araneae) as ecological indicators. Bulletin of the British Arachnological Society, 7, 83–86.

Cooper SJB, Harvey MS, Saint KM, Main BY (2011) Deep phylogeographic structuring of populations of the trapdoor spider Moggridgea tingle (Migidae) from southwestern Australia: Evidence for long-term refugia within refugia. Molecular Ecology, 20, 3219–3236.

Eggs B, Sanders D (2013) Herbivory in Spiders: The Importance of Pollen for Orb-Weavers. PLoS ONE, 8, e82637.

Feinstein J (2004) DNA sequence from butterfly frass and exuviae. Conservation Genetics, 5, 103–104.

Fetzner JW (1999) Extracting high-quality DNA from shed reptile skins: a simplified method. Biotechniques, 26, 1052–1054.

Garb JE, González A, Gillespie RG (2004) The black widow spider genus *Latrodectus* (Araneae: Theridiidae): phylogeny, biogeography, and invasion history. Molecular Phylogenetics and Evolution, 31, 1127–1142.

Gariepy TD, Lindsay R, Ogden N, Gregory TR (2012) Identifying the last supper: utility of the DNA barcode library for bloodmeal identification in ticks. Molecular Ecology Resources, 12, 646–652.

Gollan JR, Smith HM, Bulbert M, Donnelly AP, Wilkie L (2010) Using spider web types as a substitute for assessing web-building spider biodiversity and the success of habitat restoration. Biodiversity Conservation, 19, 3141–3155.

Green J (1999) Sampling method and time determines composition of spider collections. Journal of Arachnology, 24, 111–128.

Hebert PDN, Cywinska A, Ball SL, deWaard JR (2003) Biological identifications through DNA barcodes. Proceedings of the Royal Society B, 270, 313–322.

Hopkins GW, Freckleton RP (2002) Declines in the numbers of amateur and professional taxonomists: implications for conservation. Animal Conservation, 5, 245–249.

Hoy MA (1994) Insect Molecular Genetics: An Introduction to Principals and Applications. Academic Press, San Diego, California.

Huber BA (2004) The significance of copulatory structures in spider systematics. In: Biosemiotik. Praktische Anwendung und Konsequenzen für die Einzelwissenschaften (eds Schult J), pp. 89–100. VWB-Verlag, Berlin, Germany.

Huber BA and Gonzalez AP (2001) Female genital dimorphism in a spider (Araneae: Pholcidae). Journal of Zoology, 255, 301–304.

Jinbo U, Kato T, Ito M (2011) Current progress in DNA barcoding and future implications for entomology. Entomological Science, 14, 107–124.

Kremen C, Colwell RK, Erwin TL, Murphy DD, Noss RF, Sanjayan MA (1993) Terrestrial arthropod assemblages: their use in conservation planning. Conservation Biology, 7, 796–808.

Lewitus V (1935) The black widow. American Journal of Nursing, 35, 751–754.

Merrett P, Snazell R (1983) A comparison of pitfall trapping and vacuum sampling for assessing spider faunas on heathland at Ashdown Forest southeast England. Bulletin of the British Arachnological Society, 6, 1–13.

Moulder BC, Reichle DE (1972) Significance of spider predation in the energy dynamics of forest floor arthropods communities. Ecological Monographs, 42, 473–498.

Nielsen ES, Laurence AM (2000) Global diversity of insects: the problems of estimating numbers. In: Nature and human society: The quest for a sustainable world. National Academy Press, Washington DC. pp. 213–222.

Pearce JL, Venier LA (2006) The use of ground beetles (Coleoptera: Carabidae) and spiders (Araneae) as bioindicators of sustainable forest management: A review. Ecological Indicators, 6, 780–793.

Petersen SD, Mason T, Akber S, West R, White B, Wilson P (2006) Species identification of tarantulas using exuviae for international wildlife law enforcement. Conservation Genetics, 8, 497–502.

Platnick NI (2013) The world spider catalog, version 14.5. American Museum of Natural History, online at http://research.amnh.org/iz/spiders/catalog/COUNTS.html.

Riecken U (1999) Effects of short-term sampling on ecological characterization and evaluation of epigeic spider communities and their habitats for site assessment studies. Journal of Arachnology, 27, 189–195.

Robinson EA, Blagoev GA, Hebert PDN, Adamowicz SJ (2009) Prospects for using DNA barcoding to identify spiders in species-rich genera. ZooKeys, 16, 27–46.

Rosenberg DM, Danks HV, Lehmkuhl DM (1986) Importance of insects in environmental impact assessment. Environmental Management, 10, 773–783.

Salomon M (2011) The natural diet of a polyphagous predator, *Latrodectus hesperus* (Araneae: Theridiidae), over one year. Journal of Arachnology, 39, 154–160.

Samu F, Matthews GA, Lake D, Vollrath F (1992) Spider webs are efficient collectors of agrochemical spray. Pesticide Science, 36, 47–51.

Schnell IB, Thomsen PF, Wilkinson N, Rasmussen M, Jensen LRD, Willerslev E, Bertelsen MF, Gilbert MT (2012) Screening mammal biodiversity using DNA from leeches. Current Biology, 22, 262–263.

Sheppard SK, Bell J, Sunderland KD, Fenlon J, Skervin D, Symondson WOC (2005) Detection of secondary predation by PCR analyses of the gut contents of invertebrate generalist predators. Molecular Ecology, 14, 4461–4468.

Shokralla S, Spall JL, Gibson JF, Hajibabaei M (2012) Next-generation sequencing technologies for environmental DNA research. Molecular Ecology, 21, 1794–1805.

Sirvid PJ, Vink CJ, Wakelin MD, Fitzgerald BM, Hitchmough RA, Stringer IAN (2012) The conservation status of New Zealand Araneae. New Zealand Entomologist, 35, 85–90.

Sunderland KD, Fraser AM, Dixon AFG (1986) Field and laboratory studies on money spiders (Linyphiidae) as predators of cereal aphids. Journal of Applied Ecology, 23, 433–447.

Townzen JS, Brower AVZ, Judd DD (2008) Identification of mosquito bloodmeals using mitochondrial cytochrome oxidase subunit I and cytochrome b gene sequences. Medical and Veterinary Entomology, 22, 386–393.

Turner CR, Barnes MA, Xu CCY, Jones SE, Jerde CL, Lodge DM (2014) Particle size distribution and optimal capture of aqueous macrobial eDNA. Methods in Ecology and Evolution, 5, 676–684.

Vetter RS, Vincent LS, Danielsen DWR, Reinker KI, Clarke DE, Itnyre AA, Kabashima JN, Rust MK (2012) The prevalence of brown widow and black widow spiders (Araneae: Theridiidae) in Urban Southern California. Journal of Medical Entomology, 49, 947–951.

Vink CJ, Derraik JGB, Phillips CB, Sirvid PJ (2011) The invasive Australian redback spider, *Latrodectus hasseltii* Thorell 1870 (Araneae: Theridiidae): current and potential distributions, and likely impacts. Biological Invasions, 13, 1003–1019.

Zevenbergen JM, Schneider NK, Blackledge TA (2008) Fine dining or fortress? Functional shifts in spider web architecture by the western black widow *Latrodectus hesperus*. Animal Behavior, 76, 823–829.

